# ETV7 reduces inflammatory responses in breast cancer cells by repressing TNFR1/NF-κB axis

**DOI:** 10.1101/2022.09.06.506542

**Authors:** Erna Marija Meškytė, Laura Pezzè, Mattia Forcato, Irene Adelaide Bocci, Alessandra Bisio, Silvio Bicciato, Daiva Baltriukienė, Y. Ciribilli

**Author notes:** Institut für Zellbiologie, Universitätsklinikum Essen, Essen, Germany (Irene Adelaide Bocci). correspondence (YC).

## Abstract

The transcription factor ETV7 is an oncoprotein that is up-regulated in all breast cancer (BC) types. We have recently demonstrated that ETV7 promoted breast cancer progression by increasing cancer cell proliferation and stemness and was also involved in the development of chemo- and radio-resistance. However, the roles of ETV7 in breast cancer inflammation have yet to be studied. Gene ontology analysis previously performed on BC cells stably over-expressing ETV7 demonstrated that ETV7 was involved in the suppression of innate immune and inflammatory responses. To better decipher the involvement of ETV7 in these signaling pathways, in this study, we identified *TNFRSF1A*, encoding for the main receptor of TNF-α, TNFR1, as one of the genes down-regulated by ETV7. We demonstrated that ETV7 directly binds to the intron I of this gene, and we showed that the ETV7-mediated down-regulation of TNFRSF1A reduced the activation of NF-κB signaling. Furthermore, in this study, we unveiled a potential crosstalk between ETV7 and STAT3, another master regulator of inflammation. While it is known that STAT3 directly up-regulates the expression of TNFRSF1A, here we demonstrated that ETV7 reduces the ability of STAT3 to bind to the *TNFRSF1A* gene via a competitive mechanism, leading to the repression of its transcription. These results suggest that ETV7 can reduce the inflammatory responses in breast cancer through the down-regulation of TNFRSF1A.

## Introduction

Although breast cancer is becoming more curable, the cancer burden continues to increase as life expectancy rises (1). It is estimated that 1 in 8 women will develop breast cancer during their lifetime (1) and despite early diagnostic approaches and numerous available drugs, breast cancer remains the leading cause of cancer deaths in women (626,679 deaths worldwide in 2018) (2). In the vast majority of breast cancer patients, the cause of death is the development of resistance to the therapy and the development of distant metastases (3–5).

Inflammation is a hallmark of cancer and plays an important role in tumor development and progression (6); however, there are still many unknown players and mechanisms that regulate inflammatory processes in cancer. TNF-α/TNFR1/NF-κB is one of the major axis regulating inflammatory and immune processes in tumors (7). TNF-α is a pro-inflammatory cytokine that is present in the tumor microenvironment. One of its main functions is to activate NF-κB signaling by binding to tumor necrosis factor receptor 1 (TNFR1) (7). Depending on the context, this regulatory axis can promote or suppress tumor progression. On the one hand, constitutive activation of NF-κB could lead to chronic inflammation and activation of pro-tumorigenic processes such as cell proliferation, survival, invasion, and angiogenesis (8,9). On the other hand, studies have also shown that NF-κB is required for the activation of the anti-tumor immune response and a disrupted activation of NF-κB signaling may help cancer cells to escape from the host immune response (8,10–12). As there are still many unanswered questions about the effect of the TNF-α/TNFR1/NF-κB signaling pathway in cancer, it is critically important to identify transcription factors involved in the regulation of this axis.

ETV7 is a transcription factor belonging to the large family of ETS (E26 Transforming Specific) transcription factors. It is a transcriptional repressor known to be up-regulated in many cancer types (13,14). For example, ETV7 was found to be up-regulated in 85% of medulloblastoma cases and another study identified ETV7 as one of the 10 most up-regulated proteins in hepatocellular carcinoma (15,16). In 2016, Piggin and colleagues reported an increased expression of ETV7 in all types of breast cancer compared to normal breast tissue (17). Interestingly, the expression of ETV7 correlated with the aggressiveness of the tumor (17).

Previous studies showed that ETV7 promotes tumor progression by acting on various molecular and cellular pathways (14,18,19). Studies performed in our laboratory also demonstrated that the expression of ETV7 in breast cancer cells was up-regulated by different DNA-damaging agents. In the same study, we uncovered that breast cancer cells stably over-expressing ETV7 develop resistance to doxorubicin by repressing the *DNAJC15* gene, thereby increasing the expression of ABC pumps leading to the increased efflux of the doxorubicin (20). In addition, ETV7 modulates the plasticity of breast cancer stem cells, hence reducing the sensitivity of cancer cells to some anti-cancer drugs (e.g., doxorubicin, 5-fluorouracil) (19). Another study demonstrating the pro-tumorigenic activities of ETV7, reported that ETV7 can form a complex with mTOR, called mTORC3, resulting in resistance to rapamycin (an mTOR inhibitor) (21). In addition, genome-wide transcriptional profiling has demonstrated that the combined treatment of MCF7 breast cancer-derived cells with the chemotherapeutic agent doxorubicin and the inflammatory cytokine TNF-α results in the synergistic induction of ETV7 (21). ETV7 was also identified as an interferon (IFN)-stimulated gene (ISG), and its expression is known to be up-regulated upon the treatment with type I, type II, and type III interferons (22–25). We have recently shown that ETV7 negatively regulates the type I interferon response, and it is known to be involved in the viral immune response by suppressing a subset of ISGs that are important for the control of influenza and SARS-CoV-2 viruses (26,27). However, the role of ETV7 in breast cancer immunity and inflammatory processes remains to be investigated. And, since immunotherapies appear to have great potential in the treatment of solid tumors (28,29), a better understanding of the mechanisms regulating immune and inflammatory responses in breast cancer is essential for the successful development and application of novel therapeutic strategies.

In this study, we demonstrate that ETV7 plays a role in the inflammatory response in breast cancer-derived cell lines. We also show that ETV7 directly down-regulates the *TNFRSF1A* gene and reduces the activation of NF-κB signaling, thereby suppressing the inflammatory response. Moreover, we unveil a negative crosstalk between ETV7 and STAT3 in the regulation of the *TNFRSF1A* gene, and this crosstalk highlighta the importance of ETV7 in cancer immunity and inflammation. Taken collectively, we suggest ETV7 as a novel regulator of *TNFRSF1A* gene expression and, therefore, a modulator of NF-κB signaling.

## Results

### ETV7 is involved in inflammatory and immune responses

To better understand the transcriptional networks regulated by ETV7, in our previous study, we performed transcriptome analyses in two breast cancer-derived cell lines, MCF7 and T47D, that stably over-express ETV7 or empty counterpart (19). Interestingly, the most significant terms identified by gene ontology analysis of commonly down-regulated DEGs involved innate immune response and inflammatory response. Furthermore, gene set enrichment analysis (GSEA) highlighted “inflammatory response” (Fig.1A) and “TNFA_signaling_via_NF-κB” (Fig.1B) in both cell lines, confirming the involvement of ETV7 in these processes. As a result of these analyses, we obtained a list of down-regulated genes known to be involved in inflammatory pathways. For further validation, we selected a set of genes with a Fold Change lower than −1.2 in both MCF7 and T47D cell lines. Using RT-qPCR, we demonstrated the repression of all selected targets (TNFRSF1A, IL10RB, IL1R1, and TLR-2) in MCF7 and T47D (Fig. 1C) cells, with the sole exception of IL1R1, which was significantly down-regulated only in MCF7 cells.

**Figure 1.**
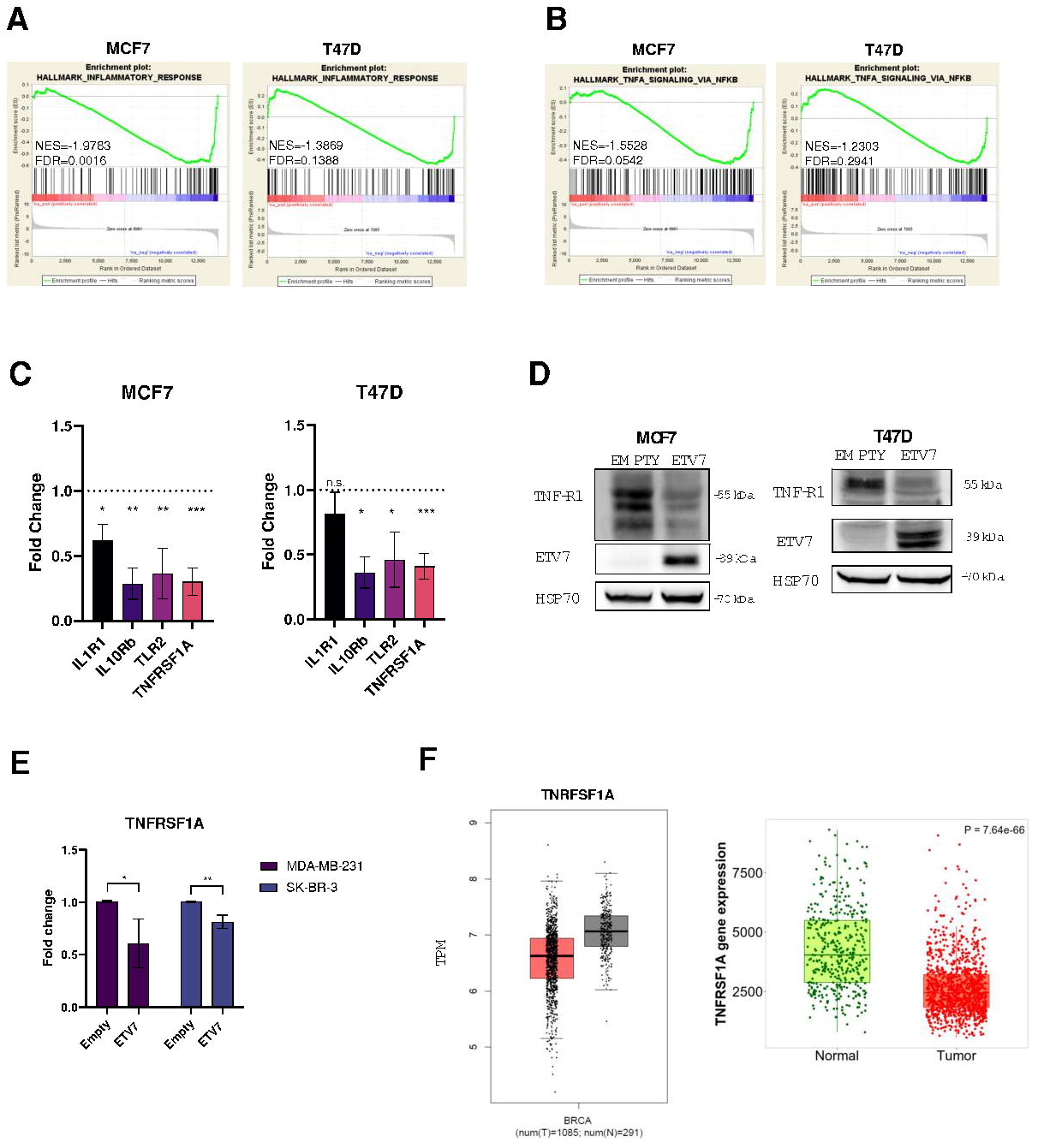
ETV7 modulates the inflammatory and immune responses in breast cancer. A-B) Gene Set Enrichment Analysis of MCF7 and T47D cells over-expressing ETV7 or its Empty counterpart. Enrichment plot for inflammatory response in MCF7 and T47D cells (A) and TNFA signalling via NFKB in MCF7 and T47D cells (B) gene sets of the Hallmark Collection. The Normalized Enrichment Score (NES) shows the degree of the enrichment of the gene set; the negative sign indicates that the gene set is down-regulated in cells over-expressing ETV7. FDR = False Discovery Rate. C) RT-qPCR analysis for the validation of genes involved in inflammation and immune response in MCF7 and T47D cells over-expressing ETV7 or Empty vector. Bars represent the averages and standard deviations of at least three biological replicates. D) The expression of TNFRSF1A at protein level in MCF7 and T47D cells over-expressing ETV7 or harbouring its empty counterpart. On the right of each blot is indicated the approximate observed molecular weight. HSP70 was used as a loading control. E) RT-qPCR analysis of the normalized expression of the TNFRSF1A gene in MDA-MB-231 and SK-BR-3 cells transiently over-expressing ETV7 or harbouring its Empty counterpart. Bars demonstrate the averages and standard deviations of at least three biological replicates. F) On the left a box plot demonstrating the differential expression analysis for the *TNFRSF1A* gene in a BRCA patients’ dataset. T= tumor (red), N=normal (grey), TPM=Transcripts per Million. GEPIA tool, based on TCGA (The Cancer Genome Atlas) databases, was used to obtain these data. On the right a box plot of TNFRSF1A expression in breast cancer patients when comparing normal and tumor RNA-seq data using TNM plot tool. Mann-Whitney test p value is shown in the right upper corner. Whole panel: * p≤0.05; ** p≤0.01; *** p≤0.001; n.s. – not significant.

### ETV7 directly down-regulates TNFRSF1A

We were particularly interested in the *TNFRSF1A* gene as it encodes for the TNFR1 receptor, which is the main receptor for TNF-α and we demonstrated that the expression of TNFRSF1A is significantly down-regulated in both MCF7 and T47D (Fig.1C) cells over-expressing ETV7. Furthermore, we were able to confirm the down-regulation of TNFRSF1A also in two other breast cancer-derived cell lines, SK-BR3 and MDA-MB-231, upon the transient over-expression of ETV7 (Fig. 1E). To understand whether this repression at the mRNA level was also reflected at the protein level, we performed Western blot analysis, demonstrating that ETV7 down-regulated TNFR1 protein in both MCF7 and T47D (Fig.1D) cells. Knowing that the increased expression of ETV7 has been detected in breast cancer patients and correlated with breast cancer aggressiveness(17), we analyzed the expression of TNFRSF1A in breast cancer patients compared to normal breast tissue. We performed gene expression analysis using the TCGA database and observed a decrease in the expression of TNFRSF1A in breast cancer tumor tissue (BRCA dataset) compared to normal tissue (Fig. 1F). Afterwards, we investigated whether the expression level of TNFRSF1A influences the survival of breast cancer patients. Using the Kaplan-Meier plotter tool, we analyzed the impact of TNFRSF1A expression in breast cancer patients and confirmed a significant correlation between lower TNFRSF1A levels and poor prognosis of breast cancer patients (Suppl. Fig. 1A).

Given the fact that ETV7 is reported to be a transcriptional repressor, to understand whether it could directly regulate the TNFRSF1A expression we searched for putative ETV7 binding sites in the regulatory elements of the *TNFRSF1A* gene. Based on ETV7 consensus sequences (30–32), we identified three potential binding sites for ETV7 containing the GGAA motif in the first intron of TNFRSF1A (Fig 2A). To understand whether the transcription repression was associated with the direct binding of ETV7 to these regulatory elements in *TNFRSF1A* gene, we performed chromatin immunoprecipitation followed by qPCR. We were able to demonstrate the direct binding of ETV7 to the TNFRSF1A intron regions (BS#1 and BS#2) in both MCF7 (Fig. 2B) and T47D (Fig. 2C) cells, hence confirming that ETV7 can directly repress TNFRSF1A.

**Figure 2.**
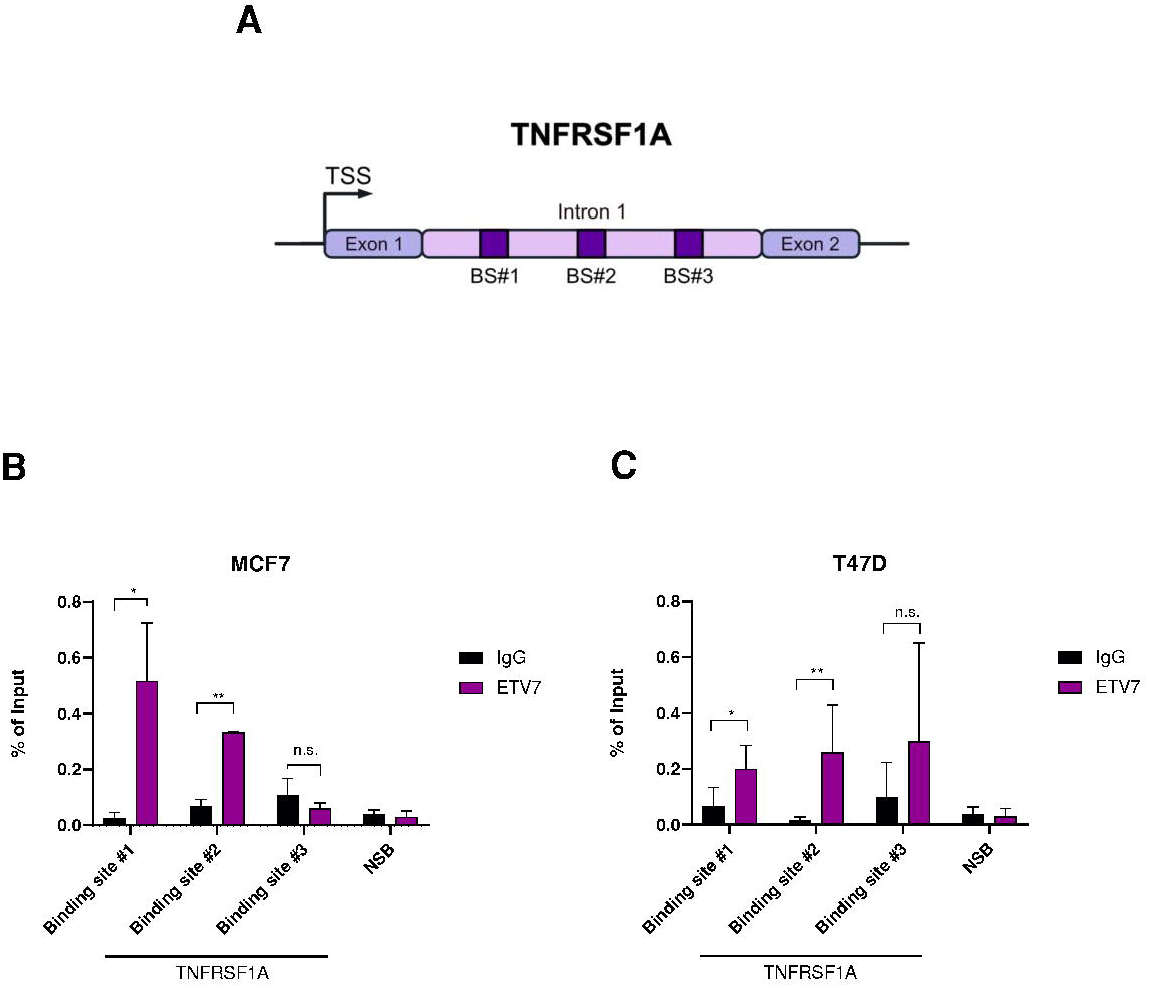
ETV7 directly represses the expression of TNFRSF1A in breast cancer cells. A) A schematic view of the TNFRSF1A Intron 1 and the studied ETV7 binding sites. TNFRSF1A BS#1 is located +5,483bp form the Transcription Start Site (TSS); BS#2 +5,627bp from TSS; BS#3 +6,069bp from TSS. B-C) ChIP-qPCR of TNSFRSF1A Intron 1 in MCF7 (B) or T47D (C) cells over-expressing ETV7. The percentage of the enrichment of ETV7 or control (normal mouse IgG) bound to TNFRSF1A Intron 1 in respect to input DNA is shown. NSB=non-specific binding, the *ACTB* promoter. Bars represent the averages and standard deviations of at least three biological replicates. * p≤0.05; ** p≤0.01; *** p≤0.001, n.s. – not significant.

### ETV7 reduces NF-κB activation by repressing TNFRSF1A

Knowing that TNF-α activates NF-κB signaling by binding to the TNFR1 receptor, we hypothesized that the ETV7-mediated repression of TNFRSF1A modulates the transcriptional activity of NF-κB in MCF7 and T47D breast cancer cells. Firstly, we performed a gene reporter assay using the pGL3-NF-κB reporter plasmid (Suppl. Fig. 1B) in MCF7 and T47D cells over-expressing ETV7 or Empty vector. Noteworthy, the over-expression of ETV7 resulted in a significant repression of the basal NF-κB transcriptional activity in both MCF7 (Fig. 3A) and T47D (Fig. 3B) cells. To understand whether this phenomenon can also be observed upon NF-κB induction, we repeated the experiment stimulating the cells with TNF-α (the main NF-κB activator), IL-6 (a broader pro-inflammatory cytokine), or a combination of these two cytokines to further stimulate the transcriptional activity. Notably, we demonstrated that ETV7-over-expressing cells did not induce the NF-κB signaling even after the stimulation (Fig. 3C and 3D). This effect was particularly strong in MCF7 cells, whereas T47D cells were overall less responsive to TNF-α stimulation. Next, we decided to investigate whether the ETV7-mediated reduction in NF-κB activation affects endogenously NF-κB target genes’ expression. We measured the mRNA expression of 4 well-known NF-κB-regulated targets, such as TNF-α, IL-8, IL-6, and A20 (33,34). The over-expression of ETV7 significantly reduced the TNF-α-dependent expression of all of these genes in both MCF7 (Fig. 3E-H) and T47D (Suppl. Fig. 1C-F) cells, with a sole exception of IL-8, whose expression was significantly reduced only in MCF7 cells. Moreover, the activation and translocation of NF-κB into the nucleus is initiated by the phosphorylation, ubiquitination, and proteolytic degradation of IκBα by the IKK complex (35,36). Therefore, we decided to investigate the effect of ETV7 over-expression on the phosphorylation of IκBα. Interestingly, the over-expression of ETV7 led to a significant reduction in phosphorylated IκBα in both MCF7 and T47D cell lines (Fig. 3I and Suppl. Fig. 1G). To confirm the hypothesis that the repression of NF-κB signaling depends, at least partially, on the ETV7-mediated repression of TNFRSF1A, we over-expressed TNFR1 in MCF7 ETV7 and MCF7 Empty cells and performed a gene reporter assay using the pGL3-NF-κB reporter plasmid (Fig. 3J). The over-expression of TNFR1 was verified in all the conditions by Western blot analysis in both MCF7 Empty and MCF7 ETV7 cells (Suppl. Fig. 1H). Notably, upon the over-expression of TNFR1, we detected a partial but significant increase in the transcriptional activity of NF-κB in MCF7 cells over-expressing ETV7 (Fig. 3K).

**Figure 3.**
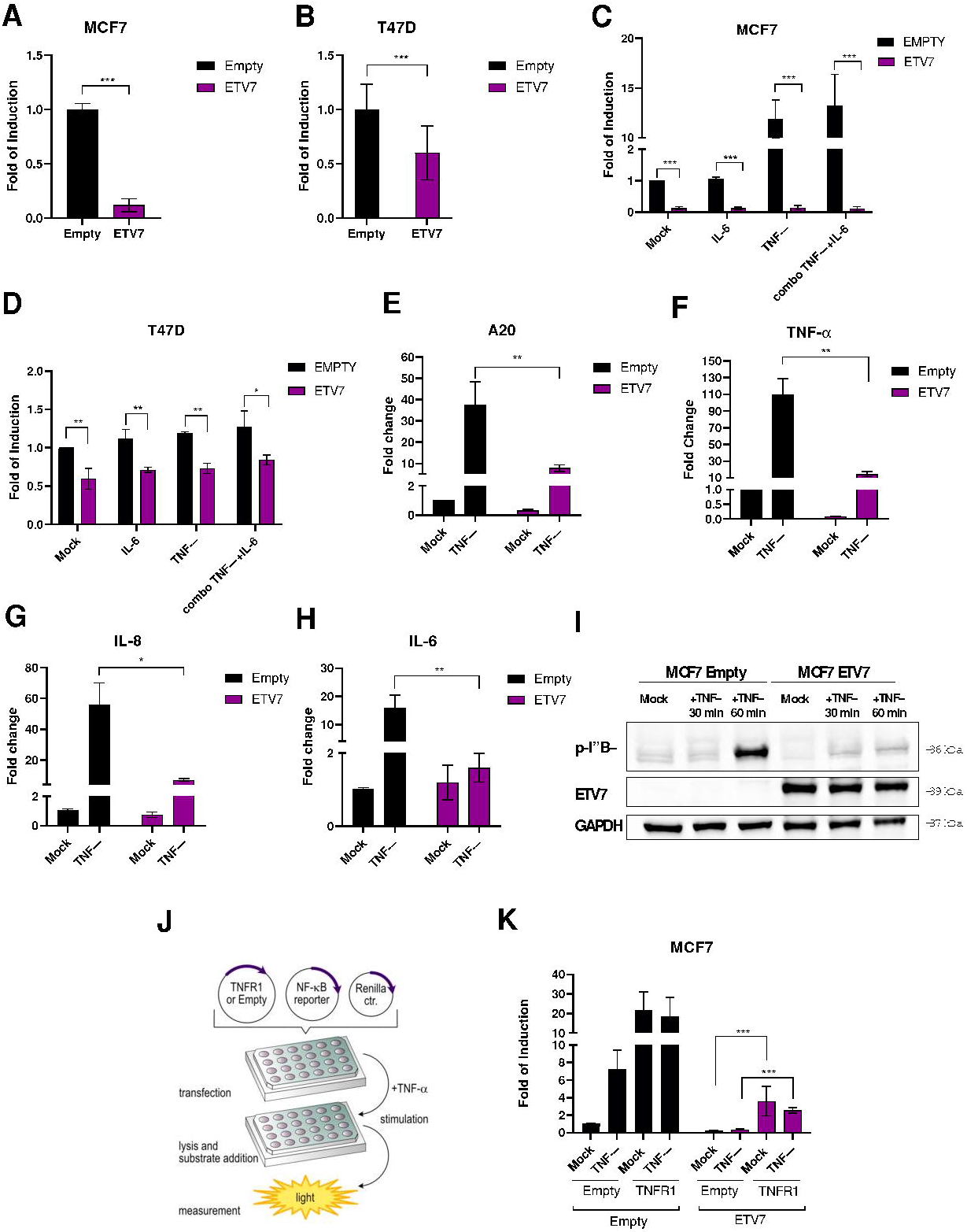
ETV7 reduces NF-κB activation by repressing TNFRSF1A. A-B) Gene reporter assays in MCF7 (A) and T47D (B) cells over-expressing ETV7, or its empty counterpart transiently transfected with pGL3-NF-κB reporter plasmid. Data is normalized using the *Renilla reniformis* luciferase reporter vector pRL-SV40 and shown as fold of induction relative to the Empty control. C-D) Gene reporter assays in MCF7 (C) and T47D (D) cells over-expressing ETV7, or its empty counterpart transfected with pGL3-NF-κB reporter plasmid and stimulated with TNF-α (10 ng/ml for MCF7 and 15 ng/ml for T47D), IL-6 (20 ng/ml) or combination of both for 4 hours. Data is normalized using the *Renilla reniformis* luciferase reporter vector pRL-SV40 and shown as fold of induction relative to the Empty control. F-I) RT-qPCR analysis of known NF-κB target genes: A20 (E), TNF-α (F), IL-8 (G) and IL-6 (H) in MCF7 ETV7 or Empty cells treated with TNF-α (10 ng/ml) for 4 hours. Bars represent the averages and standard deviations of at least three biological replicates. I) Western blot analysis of phosphorylated IκBα in MCF7 Empty and ETV7 cells in response to 10 ng/ml TNF-α treatment for 1 hours. On the right of each blot is indicated the approximate observed molecular weight. GAPDH was used as a loading control. J) A flowchart of the design of the rescue experiment. K) Gene reporter assays in MCF7 Empty or MCF7 ETV7 cells transfected with pGL3-NF-κB reporter vector and pcDNA3.1-TNFR1 or pcDNA3.1-Empty plasmids and untreated or treated with 10 ng/ml TNF-α for 4 hours. Data is normalized using the *Renilla reniformis* luciferase reporter vector pRL-SV40 and shown as fold of induction relative to the Empty untreated control. Whole panel: * p≤0.05; ** p≤0.01; *** p≤0.001.

### ETV7 competes with STAT3 in the regulation of the *TNFRSF1A* gene

It is known from the literature that STAT3 can up-regulate the expression of TNFRSF1A by binding to its first intron, the same intron bound by ETV7 for the regulation of TNFRSF1A (33). Moreover, there are similarities between the binding sites for STAT3 and ETV7 (Fig. 4A) (37,38). Hence, we decided to investigate the potential cross-talk between these two transcription factors. Not to mention that one of the binding regions (ETV7 BS #2) we identified also contains a binding site for STAT3. The potential interaction between STAT3 and ETV7 was also supported by our RNA-seq analysis, as it demonstrated a down-regulation of IL-6_JAK_STAT3 signaling in MCF7 and T47D cells (Fig. 4B and Suppl. Fig. 2A).

**Figure 4.**
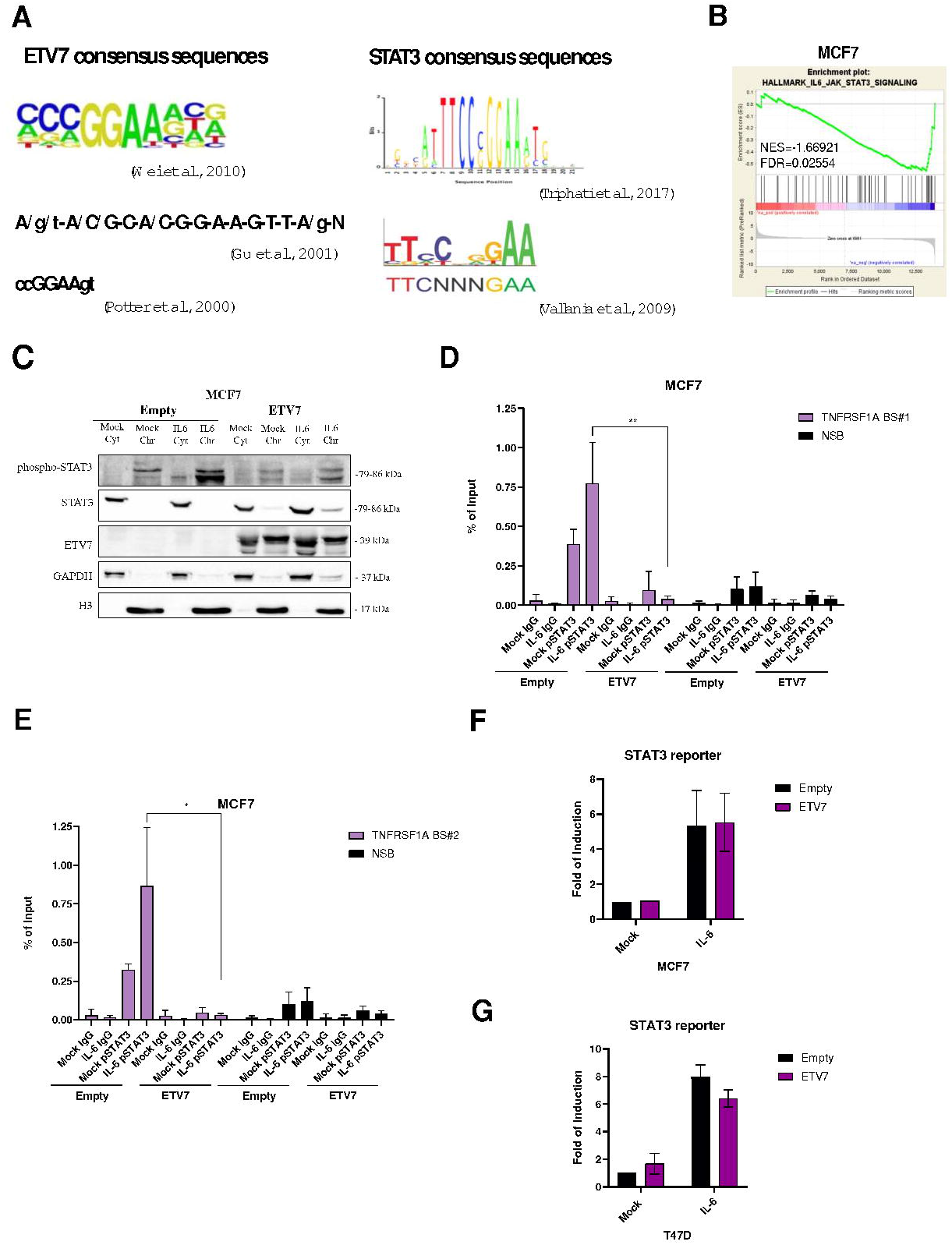
Crosstalk between ETV7 and STAT3. A) Canonical ETV7 and STAT3 binding sites known from the literature. B) Gene Set Enrichment Analysis of MCF7 cells over-expressing ETV7 or its Empty counterpart. Enrichment plot for IL6_JAK_STAT3 signalling in MCF7 cells gene sets of the Hallmark Collection. The Normalized Enrichment Score (NES) shows the degree of the enrichment of the gene set; the negative sign indicates that the gene set is down-regulated in cells over-expressing ETV7. FDR = False Discovery Rate. C) Western blot analysis of subcellular fractionation from MCF7 Empty and MCF7 ETV7 cells untreated or treated with IL-6 (20 ng/ml) for 4 hours. On the right of each blot is indicated the approximate observed molecular weight. Cyt – cytoplasmatic protein fraction, Chr – chromatin-enriched protein fraction. GAPDH was used as a loading control for the cytoplasmatic fraction. Histone H3 was used as a loading control for chromatin-enriched protein fraction. D-E) ChIP-qPCR of TNSFRSF1A Intron 1 Binding site #1 (D) and Binding site #2 (E) in MCF7 cells over-expressing ETV7 untreated or treated with IL-6 (20 ng/ml) for 4 hours. The percentage of the enrichment of pSTAT3 or control (normal rabbit IgG) bound to TNFRSF1A Intron 1 in respect to Input DNA is shown. NSB=non-specific binding, a region within the *ACTB* promoter. BS = binding site. Bars represent the averages and standard deviations of at least three biological replicates. F-G) Gene reporter assays in MCF7 Empty/ETV7 (F) and T47D Empty/ETV7 (G) cells transfected with M67-STAT3 reporter untreated or treated with IL-6 (20 ng/ml) for 4 hours. Data is normalized using the *Renilla reniformis* luciferase reporter vector pRL-SV40 and shown as fold of induction relative to the Empty untreated control. Whole panel: * p≤0.05; ** p≤0.01; *** p≤0.001.

In normal conditions, STAT3 can be activated by various external stimuli such as interferons, interleukins, and TNF-α. IL-6 is known to be one of the most potent activators of STAT3; therefore, we chose to use the stimulation with IL-6 to activate STAT3. To test whether IL-6 can also trigger the phosphorylation of STAT3 in our cellular model and to validate the subcellular localization of ETV7 and STAT3 in endogenous conditions and after the treatment with IL-6, we performed the protein subcellular fractionation followed by Western blot analysis. Data obtained confirmed that the phosphorylated form of STAT3 is mainly located in the chromatin-associated fraction and that, as expected, there is an increase in phosphorylated STAT3 in the chromatin compartment in response of the treatment with IL-6 (Fig. 4C and Suppl. Fig. 2C). ETV7 protein is located in both the cytoplasmic fraction and the chromatin-associated fraction and does not appear to be affected by the treatment with IL-6/activation of STAT3. We then analyzed the effects of STAT3 activation on the expression of the *TNFRSF1A* gene in MCF7 cells over-expressing ETV7 or an empty counterpart. Using RT-qPCR, we found that although the treatment with IL-6 induced the expression of TNFRSF1A in MCF7 empty cells, there was a significant reduction in the expression of TNFRSF1A in the cells over-expressing ETV7 (Suppl. Fig. 2B). These results confirm that ETV7 can down-regulate the expression of TNFRSF1A despite STAT3 activation.

As mentioned previously, the ETV7 binding region in TNFRSF1A is located in the first intron and it is known from the literature that STAT3 can also bind the same region (33). To understand whether ETV7 could compete with STAT3 for binding TNFRSF1A intron 1, we performed a chromatin immunoprecipitation assay. We treated ETV7 over-expressing cells with IL-6, which activates STAT3, and performed immunoprecipitation using an antibody for the phosphorylated version of STAT3. The activation of STAT3 was confirmed by performing Western blot analysis for phosphorylated STAT3, and ETV7 over-expression did not alter it (Suppl. Fig. 2E). We then analyzed by qPCR the three regions containing binding sites for ETV7 as well as STAT3. Our results demonstrated that upon the over-expression of ETV7 the ability of STAT3 to bind the 1^st^ and the 2^nd^ regulatory regions in TNFRSF1A was remarkably decreased in MCF7 cells. The tendency of the reduction in STAT3 binding to the 3^rd^ regulatory region was also visible but not statistically significant (Fig.4D-E and Suppl. Fig. 2D).

To better understand the crosstalk between these two transcription factors in the regulation of TNFRSF1A expression we tried to determine whether ETV7 could affect the of STAT3 signaling. Firstly, to verify if the over-expression of ETV7 impacted on the mRNA expression of STAT3, we analyzed STAT3 expression in MCF7 and T47D cells over-expressing ETV7 and observed that upon the over-expression of ETV7 in both cell lines, especially in MCF7, STAT3 expression was significantly down-regulated (Suppl. Fig. 2F). However, this effect was not confirmed at the protein level (Fig.4C, Suppl. Fig. 2C and 2 E). Alternatively, to evaluate the impact of ETV7 on transcriptional activity of STAT3, we performed gene reporter assays using a luciferase reporter construct containing four canonical STAT3 binding sites (4xM67 pTATA TK-Luc). Our results demonstrated that ETV7 had no effect on the overall transcriptional activity of STAT3 (Fig. 4F-G). Furthermore, we sought to understand whether ETV7 directly interacts with STAT3 protein. To evaluate this putative interaction, we performed co-immunoprecipitation (Co-IP) experiment, using antibodies against ETV7 for immunoprecipitation and detecting phosphorylated STAT3 in Western blot. According to the Co-IP results, we could not detect a direct interaction between ETV7 and STAT3 neither in MCF7 nor in T47D cells (Suppl. Fig. 2G). Overall, with this study we demonstrated the ETV7-mediated repression of TNFR1/NF-κB axis and uncovered the mechanism behind this effect depending on the competition with STAT3 in the transcriptional regulation of the TNFRSF1A gene.

## Materials and methods

### Cell lines and culture conditions

MCF7 were obtained from Interlab Cell Line Collection Bank (IRCCS Ospedale Policlinico San Martino, Genoa, Italy), and T47D cells were received from Dr. U. Pfeffer (IRCCS Ospedale Policlinico San Martino). MDA-MB-231 and SK-BR-3 cells were a gift from Prof. A. Provenzani (CIBIO Department, University of Trento, Italy).

As described previously (19), MCF7 and T47D cells were transduced with pAIP-ETV7 or pAIP-Empty lentiviral vectors to stably over-express ETV7. MCF7, SK-BR-3, and T47D were grown in DMEM medium (Gibco, ThermoFisher Scientific, Milan, Italy) supplemented with 10% of FBS (Gibco), 2mM L-Glutamine (Gibco) and a mixture of 1000U/ml Penicillin/100 μg/ml Streptomycin (Gibco); in the case of the stable over-expression with pAIP-ETV7/Empty plasmids, for the selection 0.75 μg/ml or 1.5 μg/ml of Puromycin (Gibco), respectively, was added to the medium for the selection. MDA-MB-231 cells were cultivated in DMEM medium (Gibco) supplemented with 10% of FBS (Gibco), 2mM L-Glutamine (Gibco) and a mix of 1000U/ml Penicillin/100 μg/ml Streptomycin (Gibco) and 1% of Non-Essential Amino Acids (Gibco). Cells were maintained in a humidified atmosphere at 37°C with 5% of CO_2_. Cell lines were regularly monitored for mycoplasma contamination and passed for less than 2 months after thawing.

### Cytokine Stimulations

MCF7 and T47D cells were treated with 20 ng/ml IL-6 (PeproTech, London, UK), and 10 ng/ml or 20 ng/ml TNF-α (PeproTech), respectively. Cells were stimulated for 1 hour for protein analysis and 4 hours for Chromatin ImmunoPrecipitation, luciferase reporter assay, and mRNA analyses.

### RNA extraction and RT-qPCR

Total RNA was isolated using the RNeasy Mini Kit (Qiagen, Milan, Italy) and converted into cDNA with the PrimeScriptTM RT reagent kit (Takara, Diatech Lab Line, Ancona, Italy). RT-qPCR was performed with 25 ng of the template cDNA using the qPCRBIO SyGreen 2X (PCR Biosystems, Resnova, Rome, Italy) or qPCRBIO SyGreen 2X Blue (PCR Biosystems) mastermix, and CFX384 (Biorad, Milan, Italy) was used as a detection system. YWHAZ and ACTB were used as housekeeping genes. Relative fold change was calculated using the ΔΔCt method as described previously (39,40). For primer design, the Primer-BLAST online tool (41) was used, and all primers (obtained both from Eurofins Genomics, Ebersberg, Germany and Metabion International AG, Planegg/Steinkirchen, Germany) were tested for the specificity and efficiency. Primer sequences are listed in the Supplementary Table 1.

### Western blot

Total protein cell extracts were obtained by lysing the cells with RIPA buffer supplemented with 1x protease inhibitors (PI) (Roche, Milan, Italy). Proteins were quantified using the BCA method (Pierce, ThermoFisher Scientific) and then 30-50 μg of proteins were loaded on 8-12 % polyacrylamide gels for SDS-PAGE. After the separation, the proteins were transferred on a nitrocellulose membrane (Amersham, Merck,) which was probed over-night at 4°C with specific antibodies diluted in 1-3% skimmed milk-PBS-0.1% Tween solution: TNFR1 (H-5, Santa Cruz Biotechnologies, DBA, Milan, Italy), STAT3 (124H6, Cell Signaling Technologies, Euroclone, Milan, Italy), pSTAT3 (Y705, Cell Signaling Technologies), HSP70 (C92F3A-5, Santa Cruz Biotechnologies), TEL2 (E-1, Santa Cruz Biotechnologies), GAPDH (6C5, Santa Cruz Biotechnologies), H3 (Abcam), pIKBα (Ser32/36, Santa Cruz Biotechnologies). Detection was performed with ECL Select Reagent (GE Healthcare, Cytiva) using UVITec Alliance LD2 (UVITec Cambridge, UK) imaging system.

### Cytoplasmic-nuclear fractionation

Cytosolic and chromatin-enriched protein fractions were extracted as recently described (42). Briefly, cellular pellets were resuspended in NSB buffer (10 mM HEPES, 10 mM KCl, 1.5 mM MgCl_2_, 0.34 M Sucrose, 10% Glycerol, 1 mM DTT, 0.1% TritonX100 added with 1x PI and 1x Phosphatase Inhibitors - Roche) and left on ice for 8 min. Then the samples were centrifuged at 1300 rpm at 4°C for 10 min. The supernatant, containing the cytoplasmic protein fraction, was collected. The remaining nuclei were resuspended in the NSB buffer supplemented with 1 mM CaCl_2_ and 2000 gel units/ml of MNase (New England Biolabs) and incubated at 37°C for 10 min. The MNase reaction is stopped by adding 2 mM EGTA (Sigma-Aldrich/Merck). Afterwards, the samples were centrifuged at 13000 rpm at 4°C for 10 min and the fraction containing the nuclear soluble proteins was collected. The remaining pellet was resuspended in the NSB buffer supplemented with 600 mM NaCl and incubated at 4°C overnight. Then, the samples were centrifuged at 13000 rpm at 4°C for 10 min and chromatin-enriched protein fraction was collected. The protein samples were then loaded on the SDS-PAGE gel and Western blot procedure was executed as described above.

### Plasmids and cloning

The expression plasmids pCMV6-Entry-Empty and pCMV6-Entry-ETV7 C-terminally tagged with DDK-Myc were purchased from Origene (Tema Ricerca, Bologna, Italy). pGL3-NF-κb reporter, containing *Photinus pyralis* (Firefly) luciferase gene under the control of a NF-κB responsive element was a gift from Dr. Alessio Nencioni (University of Genoa, Italy). 4xM67 pTATA TK-Luc, containing four copies of the sequence GG<u>TTCCCGTAA</u>ATGCATCA (underlined is the STAT-binding site) was obtained from Prof. David Frank (Dana-Farber Cancer Institute, Boston, CA, USA). pRL-SV40 (Promega) plasmid constituently expressing the *Renilla reniformis* luciferase cDNA was used as transfection efficiency control for gene reporter assay.

pcDNA3-TNFR1 expression vector was generated by cloning with the primers indicated below to PCR amplify (using Q5 High-Fidelity PCR kit, New England Biolabs, Euroclone, Milan, Italy) the TNFR1 reference sequence from pBMNZ-neo-Flag-TNFR1 L380A (gift from Martin Kluger, Addgene plasmid # 43949; http://n2t.net/addgene:43949) (43) and inserting it into pcDNA3.1 plasmid (the tails containing the target sequences of restriction endonucleases are indicated in lowercases):

Fw: gcggtaccATGAGGGCCTGGATCTTCTTTC

Rv: tagcggccgcTCATCTGAGAAGACTGGGCGCG

Purified PCR product was inserted into the pcDNA3.1 backbone using KpnI and NotI restriction endonucleases and T4 DNA Ligase (New England Biolabs). Correct cloning was verified by diagnostic restriction and direct sequencing (Microsynth, Balgach, Switzerland).

### Transient transfections

24 hours prior transfection, 0.2 × 10^6^ SK-BR3, MDA-MB-231, and BT549 cells were seeded in 6-well plates. Cells were transfected using Lipofectamine LTX and Plus Reagent (Life Technologies) along with 1 μg of pCMV-Entry-Empty or pCMV-Entry-ETV7 plasmid (Origene and(20)). After 48 hours the cells were collected and processed accordingly.

### Gene reporter assays

First, 90,000 cells per well were seeded in 24-well plates and after 24 hours the cells were transfected with Lipofectamine LTX and Plus Reagent (ThermoFisher Scientific) along with different combinations of the plasmids according to the experiment: 50 ng of normalizing vector pRL-SV40, 200 ng of expression vectors (pcDNA3.1-Empty/pcDNA3.1-TNFR1), and reporter vectors (350 ng 4xM67 pTATA TK-Luc and 300 ng pGL3-NF-κB). Twenty-four hours post-transfection, if necessary, the cells were stimulated with appropriate concentrations of different cytokines. Then, the cells were washed once with 1X PBS and lysed in 1X PLB (Passive Lysis Buffer) buffer (Promega). Afterwards, the activity of luciferase was measured using the Dual-Luciferase Reporter Assay System (Promega) following the manufacturer’s procedure and using the Varioskan LUX multimode microplate reader (ThermoFisher Scientific). *Renilla* luciferase activity was used as an indicator of transfection efficiency and used to obtain the Relative Light Unit (RLU) values.

### Chromatin immunoprecipitation (ChIP)

ChIP experiments were performed as previously described (20). Briefly, 3 × 10^6^ MCF7 Empty/ETV7 or T47D Empty/ETV7 cells were seeded in 15 cm dishes. The day after, if necessary, the cells were treated with 20 ng/ml IL-6 for 4 hours and afterwards cross-linked for 8 min using 1% Formaldehyde. At the end of the incubation, 125 mM Glycine was added and left for 5 min. Then, the cells were washed with ice cold 1X PBS, scraped and collected in 1X PBS supplemented with protease inhibitors (PI). Then, the pellet was lysed with lysis buffer (1% SDS) supplemented with 100 μg/ml salmon sperm single-strand DNA (ssDNA) and PI. After the lysis, the samples were centrifuged, and the supernatant was discarded. Then, the pellets were resuspended in the sonication buffer (0.25% SDS, 200 mM NaCl) supplemented with 100 μg/ml ssDNA and PI and sonicated using Bioruptor Pico sonicator (Diagenode, Denville, NJ, USA). In order to reach DNA fragments in the range of 200-700 bp, for the MCF7 cells we used 45 cycles (30 s On/ 30 s Off) and for T47D cells - 15 cycles (30 s On/ 30 s Off). Then, the samples were diluted and incubated with 2 μg the appropriate antibody or IgG (Santa Cruz Biotechnologies, mouse or rabbit according to the antibody used) and Dynabeads with protein G or A (Life Technologies) overnight at 4°C in a rotator. Input sample (10% of the sample volume) was incubated overnight at 4°C without dilution or addition of any antibodies or beads. The day after, the samples were washed through multiple washing steps and eluted at 65°C overnight by adding the elution buffer and 1X TE supplemented with 0.65% of SDS. Then the samples were processed with 50 μg of Proteinase K (ThermoFisher Scientific) for 2 hours at 56°C and 50 μg of RNase A (VWR International, Radnor, PA, USA) for 30 min, at 37°C. Afterwards, DNA was purified using QIAquick PCR purification kit (Qiagen, Germany). qPCR was performed using GoTaq^®^ qPCR Master Mix (Promega) and BioRad CFX384 qPCR system. Primer sequences are listed in the Supplementary Table 1.

### Co-Immunoprecipitation

MCF7-Empty/ETV7 or T47D-Empty/ETV7 cells were seeded in P100 dishes. After 24 hours, cells were treated with 20 ng/ml IL-6 and 4 hours post-treatment lysed using CHAPS buffer and incubated overnight with 2 μg of an anti-ETV7 antibody (TEL2, Santa Cruz Biotechnologies) or normal mouse IgG (Santa Cruz Biotechnologies) previously bound with Dynabeads protein G magnetic beads (Life Technologies). Then, the beads were washed, the immunoprecipitated lysates were eluted and loaded on a polyacrylamide gel for SDS-PAGE. Following steps were performed equally to the previously Western blot procedure.

### Gene expression profiling

The list of differentially regulated genes and enrichment scores for gene function and biological processes were obtained as we described in our recent study (19). Data were deposited with the accession number GSE152580 on the Gene Expression Omnibus database (GEO, https://www.ncbi.nlm.nih.gov/geo/). GSEA and GO results are available from our previous work (19).

### Statistics

If not indicated otherwise, statistical analyses were performed using GraphPad Prism version 9 software. For determining the statistical significance among two classes of samples, the unpaired t-test was used. Graphic illustrations were generated using Affinity designer tool (Serif, West Bridgford, UK).

### Patient databases

Data from cancer patients were obtained from available online tools; specifically, to compare the expression levels of TNFRSF1A between breast cancer and normal breast tissues, we used GEPIA (Gene Expression Profiling Interactive Analysis, http://gepia.cancer-pku.cn/, (44)) while to determine the correlation of ETV7 expression levels and prognosis in breast cancer patients we used Kaplan-Meier plotter (http://kmplot.com/analysis/, (45))

## Discussion

ETV7 is a transcriptional repressor that belongs to the family of ETS transcription factors (14). It is known to be up-regulated in many cancer types (13,14,16). In addition, Piggins and colleagues demonstrated that ETV7 expression was significantly higher in breast cancer tissues compared to normal breast tissue, suggesting that ETV7 may play an important role in breast cancer development and progression (17). Our previous studies as well as those of other scientists have already shown that ETV7 is involved in the development of drug resistance to various DNA-damaging drugs as well as mTOR inhibitor rapamycin (18–20). Furthermore, ETV7 is also well-known as an interferon (IFN)-stimulated gene. Interestingly, we recently demonstrated that ETV7 repressed a number of IFN-responsive genes, increasing the subpopulation of breast cancer stem-like cell and, thus, the resistance to chemo- and radiotherapy (19,22,23,25).

Evidences from computational studies in cancer, as well as the already known role of ETV7 in viral infections, indicate a potential function for ETV7 in cancer immunity and inflammation. However, the role of ETV7 in solid cancer tumor microenvironment, inflammation, and immune response remains to be studied (26,46). Therefore, in this study, we focused on deciphering the role of ETV7 in breast cancer immunity and inflammatory response. The RNA-seq analysis we had previously conducted on the breast cancer-derived cells MCF7 and T47D over-expressing ETV7 or its empty counterpart, supported our hypothesis by demonstrating the involvement of ETV7 in inflammation and immune responses (Fig. 1A and B). We validated several putative targets - TLR2, TNFRSF1A, IL1R1, IL10RB – that are down-regulated by ETV7 and known to be involved in inflammatory and immune processes and focused on a more in-depth analysis of the regulation of *TNFRSF1A* gene expression (47–49) (Fig. 1C).

In this study, we confirmed that the *TNFRSF1A* gene was significantly down-regulated by ETV7 at mRNA and protein levels both in MCF7 and T47D cell lines (Fig. 1C and D). We also observed the transcriptional repression of the *TNFRSF1A* gene in other breast cancer cell lines SK-BR-3 and MDA-MB-231upon transient over-expression of ETV7, as well as in the patient’s data from the TCGA (The Cancer Genome Atlas) database, which demonstrate the potential biological relevance (Fig. 1E and F).

We confirmed that the TNFRSF1A repression involves the direct binding of ETV7 to the Intron 1 of TNFRSF1A, and we identified three ETV7 binding sites in this region (Fig. 2A-C). *TNFRSF1A* encodes for Tumor Necrosis Factor Receptor 1 (TNFR1), one of the most important transmembrane receptors for TNF-α. By binding to the TNFR1 receptor, TNF-α activates NF-κB signaling, a group of transcription factors including RelA/p65, RelB, c-Rel, p50, and p52 (50–52). NF-κB is involved in the regulation of several critical cellular processes such as proliferation, cell death, survival, and cellular homeostasis (52). Furthermore, a key function of NF-κB is the control of the immune response. Indeed, NF-κB regulates the expression of different genes involved in both innate and adaptive immune responses, as well as inflammation (53,54).

Interestingly, STAT3, another master regulator of inflammation and immunity, is able to induce NF-κB activation by up-regulating TNFRSF1A (33), specifically by directly binding to its first intron, the same regulatory region also bound by ETV7 (33). However, in the breast cancer cells over-expressing ETV7 we could not observe this regulatory mechanism, even when STAT3 was activated by the treatment with IL-6 (Suppl. Fig. 2B), thus demonstrating that there is a putative competitive relationship between ETV7 and STAT3 in the regulation of the TNFRSF1A gene. Given that the consensus motifs of STAT3 and ETV7 are similar (i.e.: TTCCCGGAA and CA/CGGAAGT, respectively (30–32,37,38)), we searched for possible binding sites for ETV7 in the first intron of the *TNFRSF1A* gene that could be also used by STAT3 (Fig. 4A). After performing chromatin immunoprecipitation, we demonstrated that ETV7 is able to reduce the binding of STAT3 to the Intron 1 of TNFRSF1A, confirming our hypothesis that ETV7 competes with STAT3 when the binding sites are close to each other (Fig. 4D, E and Suppl. Fig. 2D). To further confirm the competition between ETV7 and STAT3 in the regulation of *TNFRSF1A* gene, a reporter assay could be performed in cancer cells over-expressing ETV7 using a reporter vector containing the first intron of the *TNFRSF1A* gene.

According to the literature, TNFR1 is crucial for the activation of the NF-κB signaling pathway; therefore, we aimed to determine whether the ETV7-mediated repression of the *TNFRSF1A* gene also affects the activation of NF-κB. We confirmed that in breast cancer cells over-expressing ETV7, the repression of TNFRSF1A decreased the activation of NF-κB both in the basal state and upon the stimulation with TNF-α (Fig. 3A-D). The ETV7-mediated reduction in NF-κB signaling was more pronunced in MCF7 cells compared with T47D cells, as T47D cells were globally less responsive to TNF-α. We hypothesize that this reduced sensitivity to TNF-α is due to NF-κB activity in T47D cells, which is already high prior to the stimulation. The reduced activation of NF-κB was also confirmed by the detection of the reduced level of IκBα phosphorylation in cells over-expressing ETV7 (Fig. 3I and Suppl. Fig. 1G).

Furthermore, we showed that this decreased NF-κB activation leads to a reduced expression of NF-κB target genes IL-8, IL-6, A20, and TNF-α, which are well-known to be involved in the inflammatory processes, indicating reduced inflammatory and immune processes (Fig. 3E-H and Suppl. Fig. 1C-F). However, to better characterize the biological impact of NF-κB target genes repression, the reduced expression of pro-inflammatory cytokines or chemokines should be investigated by performing secretome analysis in cells over-expressing ETV7 or the empty vector as a control. After introducing ectopic TNFR1 into our cellular systems, we confirmed that the reduced NF-κB activity was, at least partially, dependent on the ETV7-mediated repression of TNFRSF1A (Fig. 3J-K). The partial response could be explained by the fact that ETV7 also represses other elements such as Toll-like receptor 2 or IL-1 receptor 1 that are also involved in the activation of NF-κB (54,55). Collectively, our results suggest that the ETV7-mediated modulation of the TNFRSF1A expression regulates NF-κB activity in MCF7 and T47D cells. Based on our data, we propose that ETV7 represses TNFRSF1A through the displacement of STAT3 (working as positive regulator) from TNFRSF1A Intron 1 and this effect results in a significant reduction of NF-κB-dependent responses (Fig. 5).

**Figure 5.**
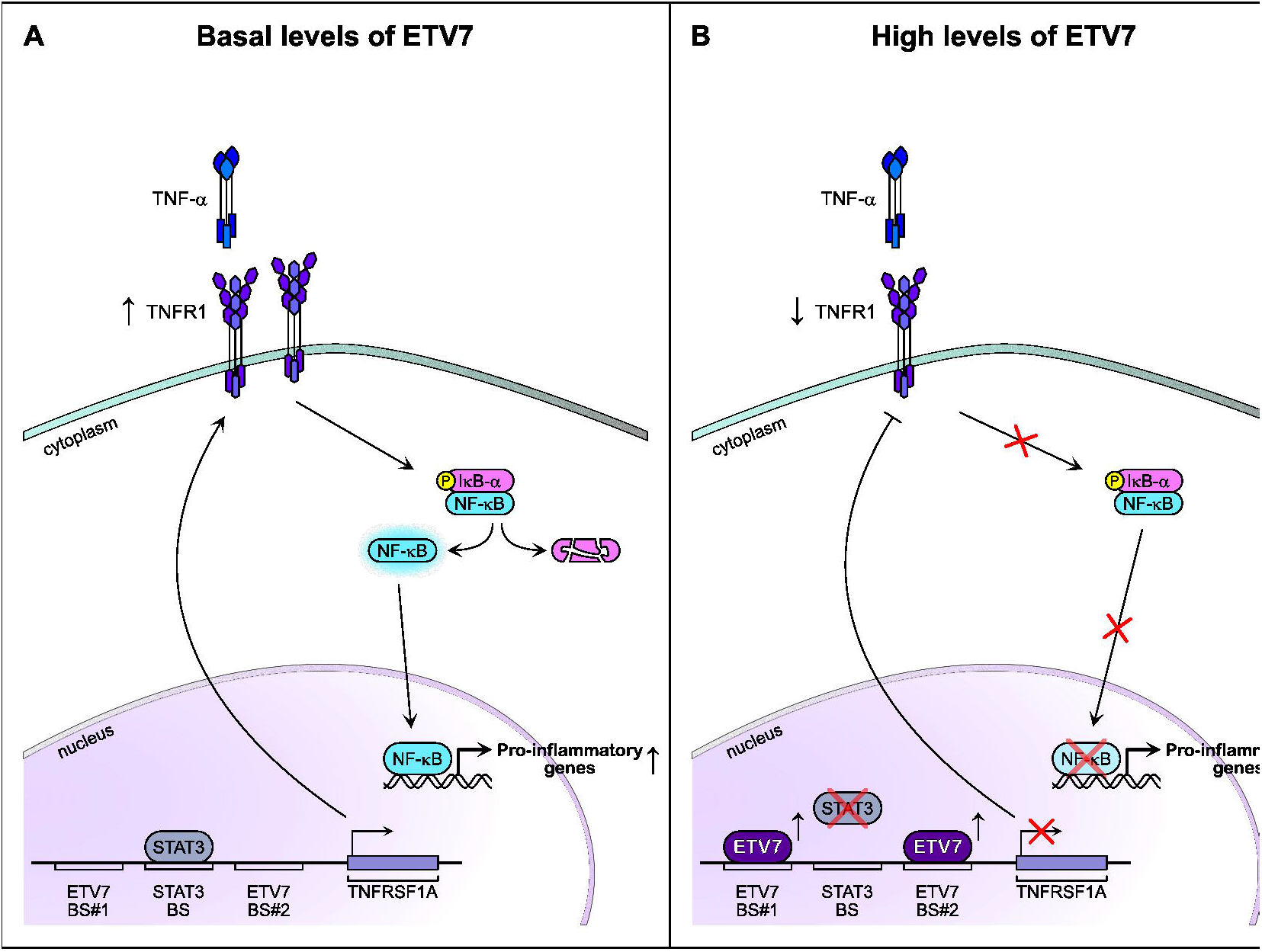
ETV7 can compete with STAT3 in the regulation of the *TNFRSF1A* gene influencing the NF-κB regulatory pathway. A) The canonical STAT3/TNF-α/NF-κB regulatory pathway. STAT3 binds to its regulatory element in the first intron of the *TNFRSF1A* gene and induce its expression. Consequently, the TNF-α receptor 1 is produced. TNF-α molecules bind TNFR1 receptor and activates NF-κB signalling pathway. B) In the context where ETV7 expression is increased, ETV7 can displace STAT3 from its binding sites on the Intron 1 of TNFRSF1A and directly repress its expression. This ETV7-mediated repression leads to the reduced activation of NF-κB signalling and, hence, reduces the expression of pro-inflammatory genes.

The role of NF-κB in tumor cells and tumor microenvironment is ambivalent and is highly dependent on the tumor context. It is widely reported that NF-κB target genes control several pro-tumorigenic processes such as proliferation, cell survival, angiogenesis, and invasion. Furthermore, NF-κB sustains the tumor-associated chronic inflammation through the production of chemokines and cytokines,. However, an increasing number of studies demonstrate the role of NF-κB as a tumor suppressor, particularly important in the regulation of anti-tumor immune response. We hypothesize that the ETV7-dependent reduced activation of NF-κB could help cancer cells evade the host immune response, as its proper stimulation is essential for both innate and adaptive immune responses (56). For example, NF-κB controls the mRNA expression and protein stability of PD-L1 in tumor cells, thereby promoting the inhibition of cytotoxic CD8^+^ T cells (57,58). Besides, a study in a pancreatic ductal carcinoma mouse model shows that TNF and TNFR1 are required for the optimal cytotoxic CD8^+^ T function and tumor rejection (59).

Furthermore, reduced activation of NF-κB leads to loss of MHC-I expression, which is one of the most important mechanisms of immune evasion in cancer (60,61). The loss of MHC-I, in turn, results in a reduced sensitivity to immunotherapy. Finally, NF-κB has many autonomous functions in immune cells in the tumor microenvironment (62,63). Even though the effect of ETV7 on the cytotoxic CD8^+^ T cells and the response to the immunotherapy remains to be studied, previous data showing the repression of IFN response in cancer and viral infections, as well as the new data demonstrated in this study strongly suggest that ETV7 may play a role in the repression of the inflammatory and immune processes in breast cancer.

Taken collectively, our present study demonstrates that ETV7 represses TNFRSF1A expression by displacing STAT3 on its regulatory element. This down-regulation leads to the reduced activation of NF-κB signaling, and thus suppresses the inflammatory and immune pathways in breast cancer cells.

## Acknowledgements

We thank Dr. U Pfeffer, Dr. A. Nencioni, Prof. D. Frank, and Prof. A Provenzani for providing cell lines and reagents, and Prof. Alberto Inga for sharing reagents and helpful discussions.

## Figure legends

**Supplementary Figure 1.** A) Kaplan-Meier curves for RFS from a breast cancer cohort according to the relative expression of TNFRSF1A obtained from KM plotter tool. The number of the patients are shown below the graph. HR (Hazardous Ratio) and the statistical analyses are reported in the right corner of the graph. B) A schematic structure of the reporter plasmid used to study the NF-κB-dependent transcriptional activity. C-F) RT-qPCR analysis of the known NF-κB target genes A20 (B), IL-6 (C), IL-8 (D), TNF-α (E) in T47D ETV7 or Empty cells treated with TNF-α (10 ng/ml) for 4 hours. Bars represent the averages and standard deviations of at least three biological replicates. G) On the left, western blot analysis of phosphorylated IκBα in T47D Empty and ETV7 cells in response to 10 ng/ml TNF-α treatment for 1 hour. On the right side of each blot is indicated the approximate observed molecular weight. HSP70 was used as a loading control. On the right, the quantification of Western blot analysis of phosphorylated IκBα in T47D Empty and ETV7 cells in response to 10 ng/ml TNF-α treatment for 1 hour. Bars represent the averages and standard deviations of at least three biological replicates. Data is normalized to the signal of HSP70, which was used as a loading control. H) Western blot analysis of the lysates obtained after performing the Luciferase assay (Fig. 3K) to control the over-expression of TNFR1 in MCF7 Empty and ETV7 cells. On the right of each blot is indicated the observed molecular weight. α-actinin was used as a loading control.

**Supplementary figure 2.** A) Gene Set Enrichment Analysis of T47D cells over-expressing ETV7 or its Empty counterpart: Shown is the Enrichment plot for IL6_JAK_STAT3 signalling in T47D cells gene sets of the Hallmark Collection. The Normalized Enrichment Score (NES) shows the degree of the enrichment of the gene set; the negative sign indicates that the gene set is down-regulated in cells over-expressing ETV7. FDR = False Discovery Rate. B) RT-qPCR analysis for the expression of TNFRSF1A after the stimulation with IL-6 (20 ng/ml) for 4 hours in MCF7 cells over-expressing ETV7 or its empty counterpart. C) Western blot analysis of subcellular fractionation from T47D Empty and T47D ETV7 cells untreated or treated with IL-6 (20 ng/ml) for4 hours. On the right of each blot is indicated the approximate observed molecular weight. Cyt – cytoplasmatic protein fraction, Chr – chromatin-enriched protein fraction. GAPDH was used as a loading control for the cytoplasmatic fraction. Histone H3 was used as a loading control for chromatin-enriched protein fraction. D) ChIP-qPCR of TNSFRSF1A Intron 1 Binding site #3 in MCF7 cells over-expressing ETV7 untreated or treated with IL-6 (20 ng/ml) for 4 hours. The percentage of the enrichment of pSTAT3 or control (normal rabbit IgG) bound to TNFRSF1A Intron 1 in respect to Input DNA is shown. NSB=non-specific binding, a region within the *ACTB* promoter. BS = binding site. Bars represent the averages and standard deviations of at least three biological replicates. E) Western blot analysis of the lysates from ChIP assay to control the phosphorylation of STAT3. F) RT-qPCR analysis of STAT3 expression in MCF7 or T47D cells over-expressing ETV7 or its Empty counterpart. G) Co-immunoprecipitation analysis of protein-protein interaction between ETV7 and pSTAT3 in MCF7 and T47D cells over-expressing ETV7. Normal IgG and INPUT were used as controls. Whole panel: * p≤0.05; ** p≤0.01; *** p≤0.001.

