## Supplementary figures and images for "ETV7 reduces inflammatory responses in breast cancer cells by repressing TNFR1/NF-κB axis"

### Supplementary Figure 1

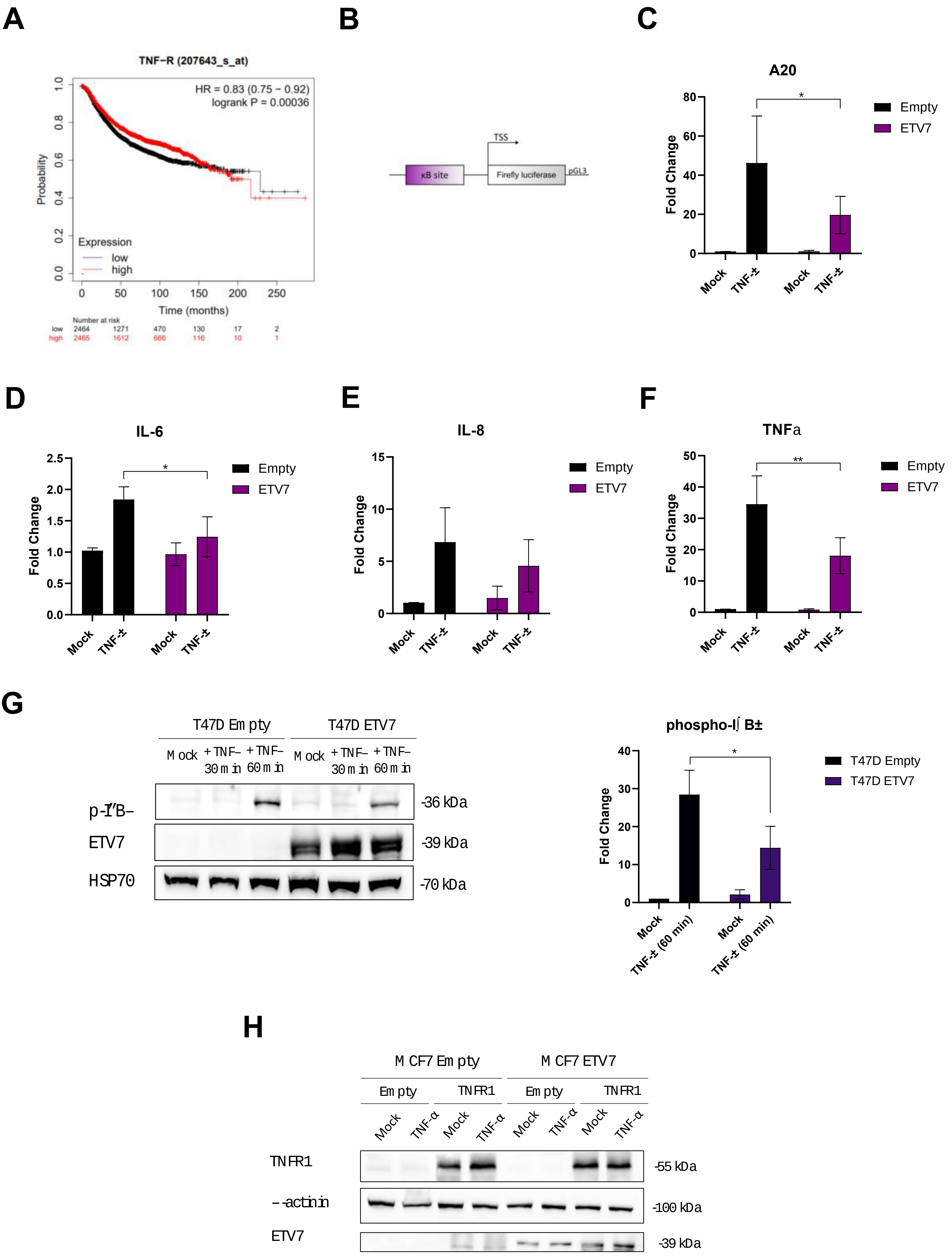

### Supplementary Figure 2

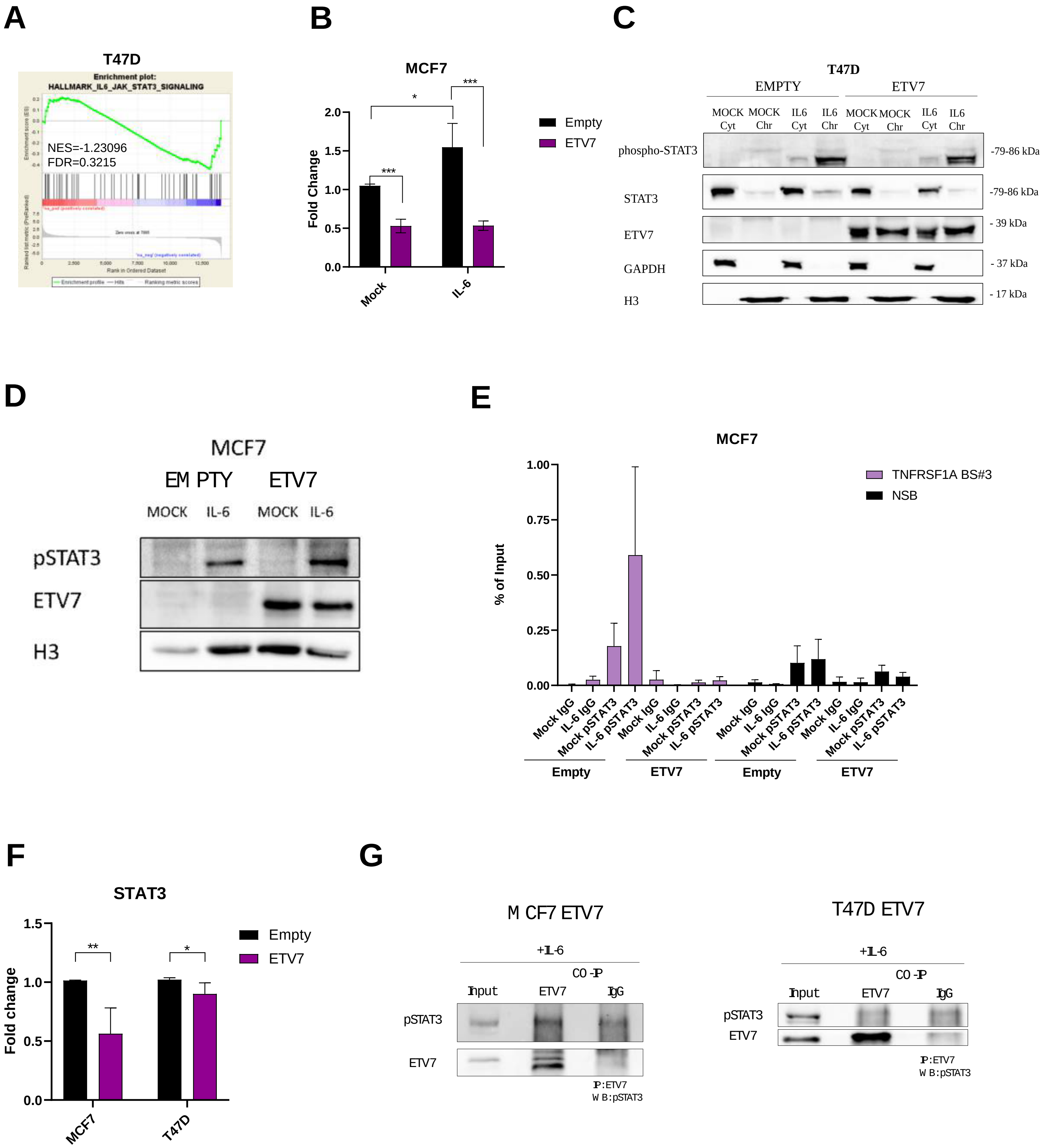
